# Cytokinetic abscission failures in a polarized epithelium affect apical membrane size and cilia

**DOI:** 10.1101/2025.09.15.675942

**Authors:** Kaela S. Lettieri, Katrina C. McNeely, Noelle D. Dwyer

## Abstract

Cytokinetic abscission is the last step of cell division, during which the intercellular bridge between daughter cells is severed. While abscission genes are linked to cancers and developmental disorders, the consequences of disrupted abscission in vivo have remained under-explored. For a polarized epithelium to expand or renew, cells within it must divide while maintaining polarity, cell junctions, and epithelial integrity. They undergo a polarized form of cytokinesis in which abscission occurs at the apical membrane. Here, we investigate how stochastic abscission failures in a polarized epithelium disrupt epithelial architecture, using mouse neuroepithelium as a model. Previously we showed that in the forebrain of *Cep55* knockout (KO) mouse embryos, a subset of neuroepithelial stem cells (NSCs) fail abscission and become binucleate, and some of those undergo p53-mediated apoptosis. Here we use the *Cep55* KO forebrain to focus on how disrupting abscission in a polarized epithelium affects the apical membrane structure. We find that NSCs in *Cep55* KO neuroepithelium have preserved epithelial polarity and integrity. However, they have enlarged apical membranes (called apical endfeet), longer primary cilia, and increased bi-ciliation. We test whether the enlarged apical endfeet arise from filling the space of apoptotic neighbors. However, blocking apoptosis does not rescue but exacerbates the phenotypes: extra-large apical endfeet have further increased multi-ciliation, supernumerary centrosomes, and abnormal or multiple nuclei. These findings show the importance of proper abscission in a polarized epithelium to maintain epithelial structure, and the need for p53-mediated apoptosis to protect the tissue in the face of stochastic abscission failures.

## INTRODUCTION

Cytokinetic abscission, the last step of cell division, is critical for proper organism development. After mitosis, the two daughter cells are connected by an intercellular bridge containing an organelle called the midbody (MB). The midbody contains dense antiparallel microtubules as well as recruits and scaffolds conserved abscission protein machinery to sever the connection between the two daughter cells (Andrade & Echard, 2022). Despite numerous studies in cell lines and single-cell systems uncovering the basic mechanisms of cytokinetic abscission, the consequences of disrupted abscission in developing tissues such as epithelia have remained under-explored. Here, we investigate how stochastic abscission failures in a polarized epithelium disrupt epithelial architecture, using the mouse neuroepithelium as a model.

The neuroepithelial stem cells (NSCs) of the embryonic cerebral cortex reside in a pseudostratified epithelium and exhibit apicobasal polarity (**Figure 1A-B’**) (McNeely & Dwyer, 2021; Norden, 2017). The apical membranes of these NSCs are called apical “endfeet” due to their highly elongated morphology. The apical endfeet, which are connected by adherens junctions and form the surface of the lateral ventricles, each contain a primary cilium that protrudes into the ventricle. Each primary cilium is anchored to the cell membrane by the basal body, a modified centrosome that templates the cilium. Additionally, during the cell cycle, the nuclei of NSCs move up and down within the cells. When an NSC divides, it must split the apical membrane between its daughters, so the nucleus must migrate to the apical membrane to undergo mitosis and cytokinesis. In these cells, as in other polarized epithelia, cytokinesis is polarized such that the cleavage furrow ingresses towards the apical membrane, forming the MB at the ventricle surface. Once the NSCs have completed telophase, the nuclei migrate back basally during G1 phase, leaving the midbody at the apical membrane to complete abscission (McNeely & Dwyer, 2020).

**Figure 1.**
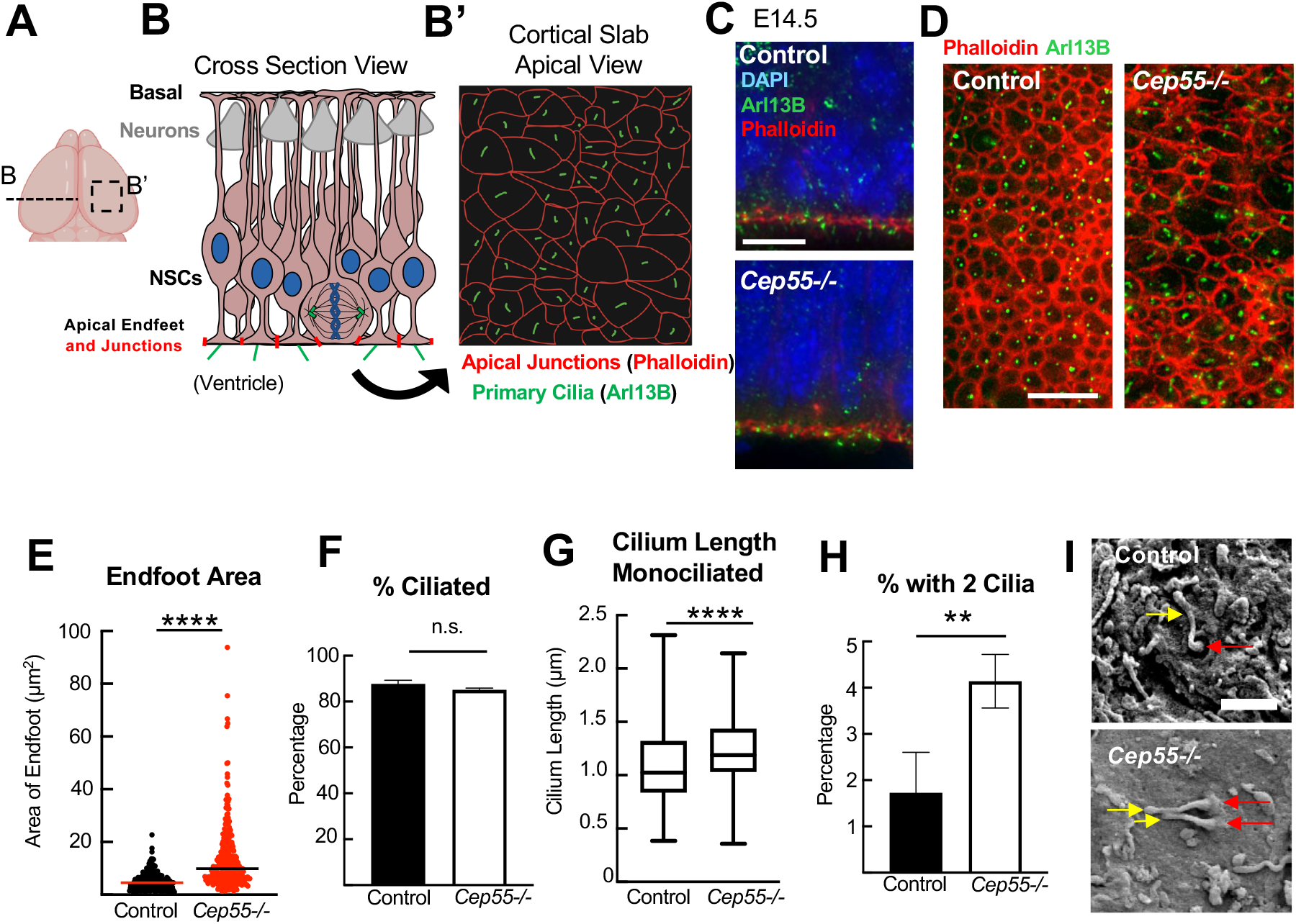
*Cep55* KO neuroepithelial stem cells have enlarged apical membranes and primary cilia phenotypes. **(A-B’)** Schematics of E14.5 mouse cerebral cortex, cross-section and apical membrane views. Neural stem cells (NSCs) form a polarized pseudostratified epithelium. A, B’ made with Biorender. **(C)** Cortical cross-sections stained with phalloidin and cilia marker Arl13b show preserved polarization in *Cep55* ko(-/-) neuroepithelium. **(D)** *En face* view of cortical slab apical membrane labeled with phalloidin and Arl13b shows enlarged apical endfeet in *Cep55* ko brains. **(E)** Median apical endfoot area is significantly increased in *Cep55* ko NSCs. **(F)** Average percentage of ciliated apical endfeet is similar in control and *Cep55* ko. **(G)** Median cilium length is slightly increased in the *Cep55* ko. **(H)** Percentage of bi-ciliated endfeet is increased in the *Cep55* ko brains. **(I)** Scanning electron microscopy (SEM) of apical membrane of cortex showing single and bi-ciliated endfeet in control and *Cep55* KO. Red arrows indicate base; yellow arrows indicate tip. * p < 0.05, ** p < 0.01, **** p < 0.001; n.s. not significant. Kolomogorov-Smirnov (K.S.) and Mann Whitney (M.W.) tests for E, G; t-test for F, H. For F, H, n=1 +/+, 2 +/−brains (2292 control endfeet) and 5 *Cep55* −/− brains (2270 endfeet). For E n=174 endfeet (1 +/+, 2 +/−brains) and n=300 *Cep55* −/− endfeet (5 brains). For G, n= 150 control endfeet (1 +/+, 2 +/− brains) and 250 *Cep55* −/− endfeet (5 brains). Scale bars 10µm in C, D; 1µm in I.

Cep55 is a key abscission regulator that specifically localizes to the late-stage midbody, and recruits and scaffolds downstream abscission protein machinery (Elia et al., 2011; Little & Dwyer, 2021; Zhao et al., 2006). We and others previously found that deletion of *Cep55* in mice resulted in delays and stochastic failures of abscission in embryonic brain NSCs, leading some to become binucleate, or undergo p53-dependent apoptosis. The loss of NSCs caused a small brain phenotype (microcephaly). However, other organs were normal size, and even in the brain, most NSCs completed abscission successfully (Little et al., 2021; Tedeschi et al., 2020).

Here we used the *Cep55* deletion (KO) mouse brain to focus on how disrupting abscission in a polarized epithelium affects apical membrane structures. We find that the *Cep55* KO neuroepithelial stem cells have longer primary cilia, increased bi-ciliation, and enlarged apical endfeet. We tested whether the enlarged apical endfeet were caused by a subset of NSCs undergoing apoptosis, leaving space for neighbors to expand. However, blocking apoptosis exacerbates the enlargement of apical endfeet rather than rescuing it. The resulting extra-large endfeet have increased multi-ciliation and abnormal or multiple nuclei. Overall, this study emphasizes the importance of proper cytokinetic abscission even in epithelia where the cells remain attached, and the importance of programmed cell death to maintain epithelial structure in the face of stochastic abscission failures.

## RESULTS

### Neuroepithelial stem cells (NSCs) in *Cep55* KO embryos have larger apical endfeet, increased bi-ciliated endfeet, and longer primary cilia than in controls

To begin assessing the how failures in cytokinetic abscission affect neuroepithelial structure, we used immunohistochemistry on embryonic cerebral cortex (**Figure 1A-D**). Fixed E14.5 cortices were stained with phalloidin to mark the apical junctions, which are enriched in F-actin, and Arl13B antibody to label primary cilia (Larkins et al., 2011). Staining these markers on cross-sections shows a bright line of actin signal and primary cilia along the apical membrane/ventricular surface, in both control and knockout (KO) sections, demonstrating preserved apico-basal polarity (**Figure 1B, C**). To image the apical surface *en face*, we immunostained fixed whole-mounts of dissected neocortex, which we call “cortical slabs” (Janisch & Dwyer, 2016). By this apical view, the apical endfeet appear as a sheet of polygons outlined by actin, with a primary cilium at the center (**Figure 1B’, D**). In *Cep55* KO images, some apical endfeet appear larger (**Figure 1D**). Indeed, by measuring the areas of individual endfeet, we found that the median apical endfoot area is about twice as large (~10µm^2^) in the *Cep55* KO neuroepithelia as controls (~5µm^2^), and the range of sizes is much wider (**Figure 1E**).

Most NSCs in the developing cerebral cortex have a primary cilium on their apical endfoot, protruding into the lateral ventricle. The percentage of ciliated endfeet (~86%) is similar in the controls and *Cep55* KOs (**Figure 1F**). However, the cilia of KO NSCs are slightly longer than controls (**Figure 1G**). Furthermore, while bi-ciliated endfeet are rare in control brains, they are increased 2-fold in the *Cep55* KO, from 2% to 4% (**Figure 1H**). Scanning electron microscopy (SEM) of cortical slab apical surfaces occasionally captured what appeared to be a bi-ciliated cell on the *Cep55* KO neuroepithelium (**Figure 1I**).

We next questioned whether the enlarged apical endfeet are related to the ciliation phenotypes. First, we wondered whether cells with 2 cilia have larger endfeet. By comparing the apical areas of endfeet having 1 versus 2 cilia, we found that control areas are similar regardless of whether they are mono-or bi-ciliated (medians ~4.2µm^2^ vs 6µm^2^) as are KO endfeet (medians ~9µm^2^ vs ~13.5µm^2^)(Dunn’s multiple comparisons). Nonetheless, both mono- and bi-ciliated *Cep55* KO endfeet are significantly larger than controls (**Figure 2A, B**). Strikingly, many of the apical endfeet areas in the *Cep55* KOs are beyond the range of control endfeet. We call these “Extra Large (XL)” endfeet (>16µm^2^). While 23% of the mono-ciliated KO endfeet are XL, the XL percentage is almost doubled, 45%, in the bi-ciliated KO endfeet. By contrast, only 6% of control bi-ciliated endfeet are XL (**Figure 2C**). Next, we asked if there was a relationship between primary cilium length and apical endfoot area. Indeed, a Pearson correlation test confirms a significant positive correlation between these measurements in the *Cep55* KOs regardless of whether they have one or two primary cilia. However, they are not correlated in controls (**Figure 2D, E**). Together, the data in Figures 1 and 2 show that the loss of Cep55 in the neuroepithelium has a big effect on apical endfoot areas, and a smaller effect on bi-ciliation and cilium length. Further, these effects are correlated within individual endfeet of the KO brains.

**Figure 2.**
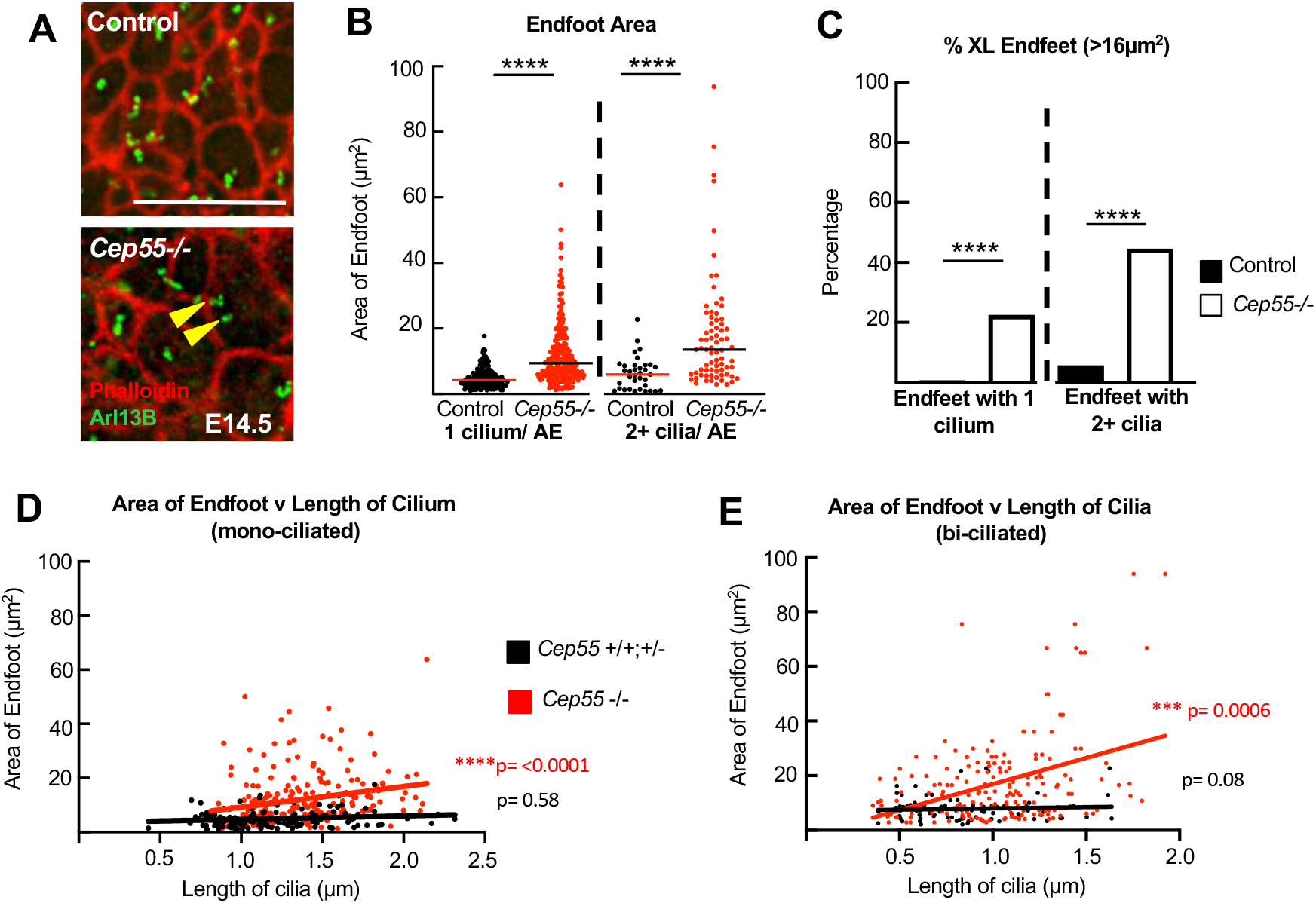
Enlarged apical endfoot areas correlate with bi-ciliation and cilium length in *Cep55* ko neuroepithelium. **(A)** Apical view of E14.5 cortical slabs labeled with phalloidin (apical junctions) and anti-Arl13b (cilia) show most NSC’s have an apical endfoot with a single cilium, but some have two cilia (yellow arrowheads). **(B)** Mono-ciliated and bi-ciliated *Cep55* KO apical endfoot areas are both significantly larger than controls. **(C)** Extra-large (XL) endfeet (area >16um^2^) are rare in control neuroepithelium, but significantly increased in *Cep55* KO, and further increased among bi-ciliated KO endfeet. **(D**,**E)** Positive correlation between individual endfeet areas and length of their cilium in *Cep55 ko*, in endfeet with one cilium (D) or two or more cilia (E). **** p < 0.001, Kolomogorov-Smirnov (K.S.) and Mann Whitney (M.W.) tests for B; Fisher’s Exact test for C; Pearson Correlation test for D, E. n=174 control endfeet (1 +/+, 2 +/− brains) and n= 300 *Cep55* −/− endfeet (5 brains). Scale bar 10µm ***Cep55 −/− p53 −/−***

### Blocking apoptosis exacerbates enlarged endfoot phenotype and results in supernumerary primary cilia and centrosomes in *Cep55;p53* dKOs

The results above raise the question of how these phenotypes arise in the *Cep55* KO brains, and if they are secondary to the stochastic abscission failures in the epithelium. We showed previously that apoptosis of NSCs is much higher in embryonic *Cep55* KO cortices than control cortices (Little et al., 2021). Therefore, we hypothesized that when NSCs undergo apoptosis and delaminate their apical attachments, the surviving NSCs expand their apical endfeet to fill the space. To test this idea, we blocked apoptosis in the *Cep55* KO embryos to determine whether that rescued the enlarged apical endfoot phenotype. Since we had shown that the apoptosis in the *Cep55* KO brains is p53-dependent (Little et al., 2021), we prevented apoptosis by creating double KOs (dKOs) of *Cep55* and *p53* (gene *Trp53)*. The *p53* KO itself has normal cortical development (Insolera et al., 2014; Marthiens et al., 2013), so we used *p53* single KOs as littermate controls.

When we examined the apical membranes of the *Cep55;p53* dKO embryo brains, we found that the enlarged endfoot phenotype was not rescued, but instead exacerbated, with even larger endfoot areas than in the *Cep55* KO (**Figure 3A, B; compare to 2A**,**B**). Separating out the mono- and multi-ciliated XL endfeet, we found that 2% of the *p53* KO mono-ciliated endfeet are XL compared to 19% in the dKOs. This percentage is doubled among the multi-ciliated endfeet, with 4% XL endfeet in the *p53* KO and 38% in the dKO (**Figure 3C**). These data demonstrate that the enlarged apical endfoot phenotype of the *Cep55* KO is not a consequence of filling space left by adjacent apoptotic neighbors.

**Figure 3.**
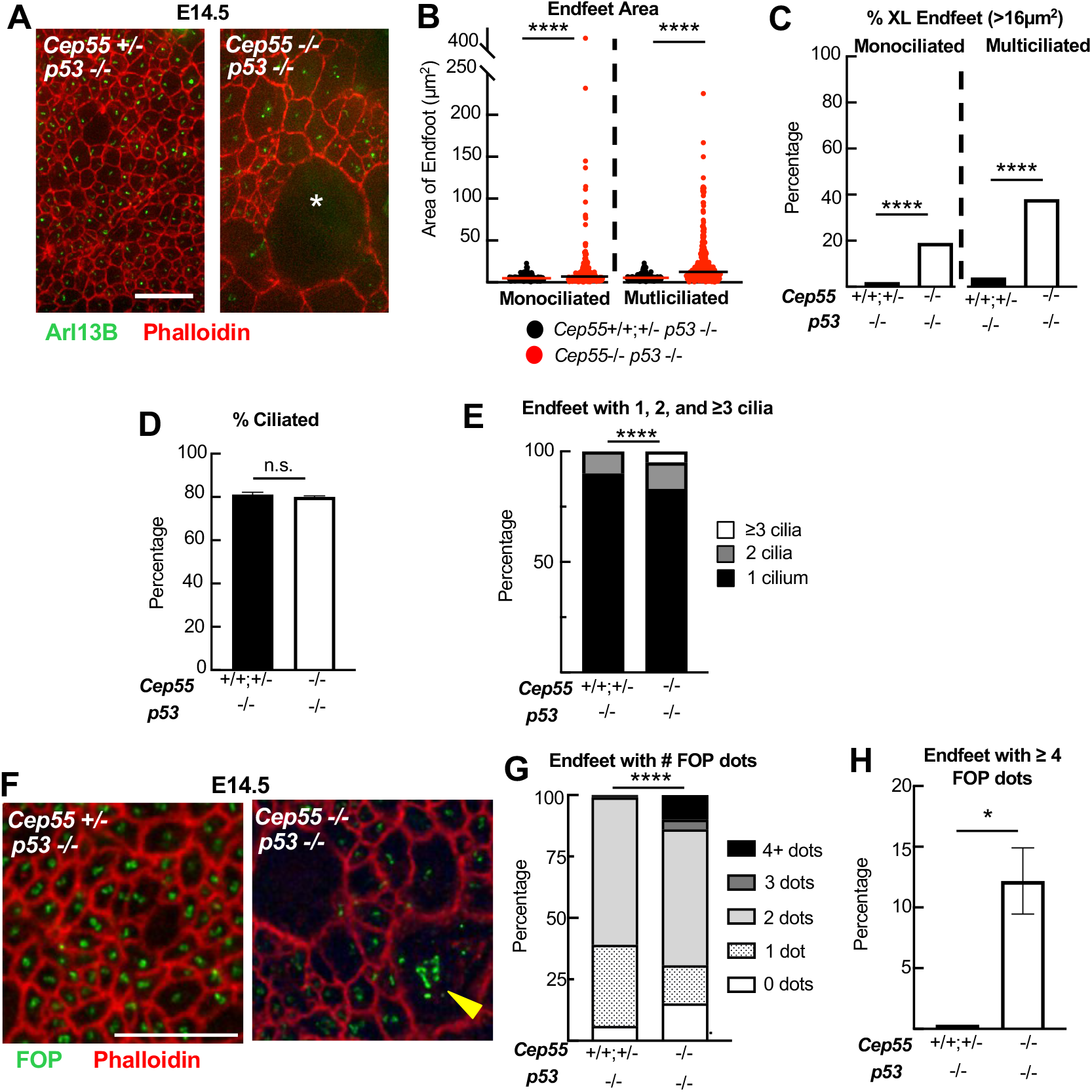
Blocking apoptosis does not rescue but exacerbates apical endfoot enlargement and results in supernumerary primary cilia and centrosomes in *Cep55;p53* double KOs. **(A)** Apical views of E14.5 cortical slabs stained with phalloidin (apical junctions) and Arl13b (cilia) show severely enlarged and irregular apical endfeet in *Cep55;p53* dKO, in which apoptosis is prevented. Asterisk indicates extra-large endfoot. **(B)** Apical endfoot areas are significantly increased in the *Cep55;p53* dKO over p53ko control. The y-axis range is extended compared to the single Cep55 ko in Fig 2B. **(C)** Percentage of XL endfeet is significantly increased in dKOs compared to *p53* KO controls. **(D)** Average percentage of ciliated apical endfeet is similar. **(E)** Proportions of apical endfeet containing 2 or 3 cilia are significantly increased in *Cep55;p53* double KO compared to *p53* single KO control. **(F)** Apical view of E14.5 cortical slabs stained with anti-FOP (centrosomes) and phalloidin. Yellow arrowhead indicates an apical endfoot with extra FOP dots in *Cep55;p53* dKO. **(G)** Proportions of apical endfeet with 0, 1, 2, 3, or ≥4 dots of FOP are significantly different when *p53* and *Cep55* are both deleted. **(H)** Percentage of apical endfeet with ≥4 FOP dots is significantly increased in the dKO compared to *p53* KO controls. * p < 0.05; ** p < 0.01, *** p <0.001, **** p <0.0001; n.s. not significant. Kolmogorov-Smirnov (K.S.) and Mann-Whitney (M.W.) tests for B, E; Fisher’s Exact test for C; t-test for D, I; Chi Square test for F, H. For B, C, *p53* KO n=312 endfeet (3 brains); *Cep55;p53* dKO n=605 endfeet (4 brains). For D, *p53* KO n=3 brains (2038 endfeet), *Cep55;p53* double KO n=4 brains (2351 endfeet). For E, *p53* KO n=1656 endfeet (3 brains), *Cep55;p53* double KO n=1875 endfeet (4 brains). For G, *p53* single KO n= 3174 endfeet (2 brains), *Cep55;p53* dKO n=1805 endfeet (3 brains). For H *p53* single KO n=2 brains (1656 endfeet), *Cep55;p53* double KO n= 4 brains (1875 endfeet). Scale bars 10µm.

An alternative possibility is that the enlarged apical endfeet in the *Cep55* KO neuroepithelium represent binucleate NSCs arising from failed cytokinetic abscission events, and preventing apoptosis increases the occurrence of enlarged binucleate cells. Consistent with this idea, we observed an increase in bi-ciliated endfeet in the *Cep55* KOs (**Figure 1H**), which could occur in cells with two nuclei and two centrosomes/basal bodies. To corroborate this theory, we examined the primary cilia in the *Cep55;p53* dKOs. Even though the percentage of ciliated cells is unchanged between the control and dKO, we indeed observed that the dKOs have triple the number of bi-ciliated endfeet compared to the single *Cep55* KOs (12% vs 4%). Furthermore, 5% of the dKO endfeet contain three or more cilia (**Figure 3D, E**).

We next questioned whether the multi-ciliated cells in the dKOs also contained extra centrosomes since centrosomes form the basal bodies of primary cilia at the plasma membrane. We immunostained for centrosome marker FOP (FGFR1 Oncogene Partner), which can appear as 1 or 2 dots, depending on distance between the centrioles and cell cycle phase (Acquaviva et al., 2009). We found that in control neuroepithelium, the majority of endfeet have 2 FOP dots and about a third have 1 dot (**Figure 3F**). In *Cep55;53* dKO neuroepithelium, more endfeet have 3 or 4 dots, with ~12% of the dKO endfeet containing ≥4 FOP dots compared to almost none in the *p53* KO controls (**Figure 3F-H**). Together, these data lead us to speculate that in the *Cep55;p53* dKOs, the NSCs that fail abscission are unable to undergo apoptosis, thus generating a larger population of bi- and multi-nucleate cells with XL endfeet, multiple primary cilia, and supernumerary centrosomes.

### XL apical endfeet in the *Cep55;p53* dKOs have abnormal and multiple nuclei

To test whether the XL and multi-ciliated endfeet observed in the *Cep55;p53* dKOs were due to multi-nucleate NSCs, we imaged the nuclei at the apical membrane within XL endfeet. Immediately, we noticed that both the non-mitotic and mitotic nuclei in the dKO appeared abnormally shaped and larger than normal (**Figure 4A**). Indeed, the nuclear area of both the mitotic and non-mitotic nuclei are significantly increased in the XL endfeet of the dKOs compared to XL endfeet of controls (**Figure 4B**,**C**). We speculate that the enlarged nuclei within the XL endfeet may form when nuclei fuse. Furthermore, some XL endfeet in the dKOs had more than one nucleus. While all control XL endfeet contained one nucleus, 25% of the dKO XL endfeet contained two nuclei and 11% contained 3 or more nuclei (**Figure 4D, E**). Strikingly, we observed that some XL endfeet contain not only multiple nuclei, but a mix of both mitotic and non-mitotic nuclei (**Figure 4F**). Finally, we examined whether the multinucleate dKO XL endfeet also have multiple primary cilia. Indeed, almost half (46%) of multinucleate XL endfeet have ≥2 primary cilia, compared to a third (32%) of the mononucleate XL endfeet in the dKO (**Figure 4G**). Together these data support the hypothesis that the extra-large apical endfoot areas and multi-ciliation arise from NSCs that fail abscission, escape apoptosis, and form bi- or multi-nucleate cells.

**Figure 4.**
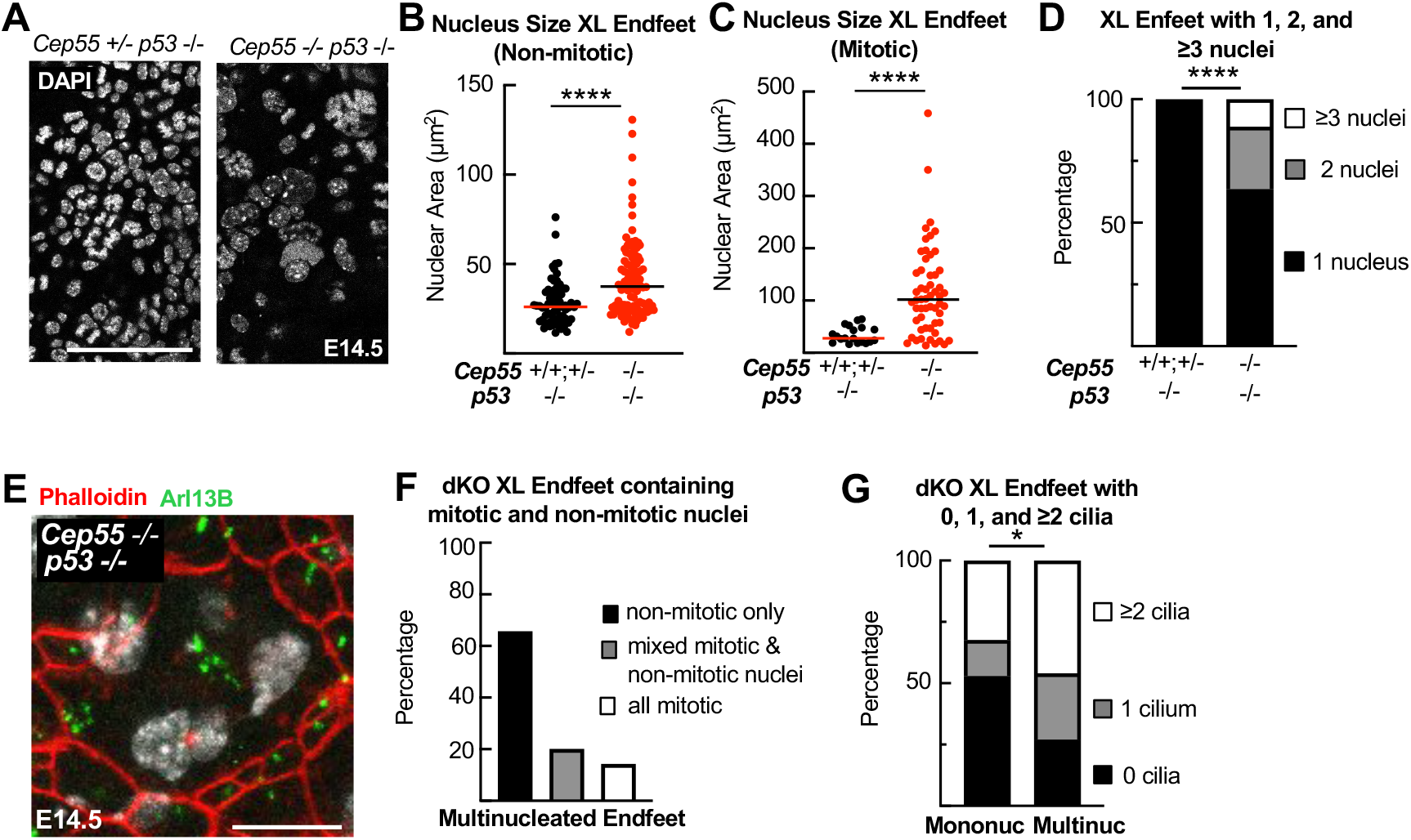
Extra-large apical endfeet in the *Cep55;p53* dKO contain abnormal and multiple nuclei. **(A)** Images of nuclei near the apical surfaces of E14.5 cortical slabs from *p53*−/− control and *Cep55;p53* dKO embryos show several large irregular nuclei in *Cep55;p53* dKO. **(B, C)** Nuclei in XL (extra-large) endfeet are bigger in *Cep55;p53* dKO compared to controls, whether non-mitotic or mitotic (by chromatin appearance). **(D)** Proportions of XL endfeet containing 1, 2, or ≥3 nuclei. Multinucleated endfeet are increased in *Cep55;p53* dKO versus *p53 −/−*control. **(E)** Apical view image of an XL apical endfoot in *Cep55;p53* dKO containing multiple nuclei and cilia (Arl13b marks cilia in green; DAPI marks nuclei in white; phalloidin marks apical junction actin in red). **(F)** Multinucleated XL endfeet in *Cep55;p53* dKO neuroepithelia usually contain only non-mitotic nuclei, but can contain mitotic or mixed nuclei. **G)** Multinucleate XL endfeet in *Cep55;p53* dKO are more likely to have ≥2 cilia compared to mononucleate XL endfeet. *p<0.05, ****p< 0.0001; Kolmogorov-Smirnov (K.S.) and Mann-Whitney (M.W.) tests for B, C; Chi square for D, G. n= 84 endfeet for *p53* single KO (2 brains); n= 99 endfeet for dKO (4 brains). Scale bars 50µm for A, 10µm for E

### Discussion

Polarized epithelia have a specialized form of cytokinesis, in which there is not a contractile ring, but a contractile arch making an asymmetric cleavage furrow in the basolateral membrane; then, midbodies form and mediate abscission at the apical membrane. *A priori*, one might think that failure to complete abscission would not cause a problem since the cells are already attached to each other via apical junctions. However, our previous work showed how disruptions of abscission in embryonic neuroepithelium profoundly reduce neurogenesis and brain size, partly by causing p53-mediated apoptosis in some cells (Janisch et al., 2013; Little & Dwyer, 2019;Little et al., 2021). Here we focused on how abscission failures affect the neuroepithelial membrane structure. We used *Cep55* KO mice as a tool to cause stochastic abscission failures and test how it perturbs the neuroepithelium of the embryonic forebrain. We found that the apical endfeet of KO neuroepithelial cells have significantly larger areas. In addition, their primary cilia have increased lengths, correlating with their apical membrane area. These phenotypes are not due to apoptosis because blocking apoptosis exacerbated these phenotypes, resulting in extra-large, multi-ciliated apical endfeet, with supernumerary centrosomes and abnormal or multiple nuclei. It appears that when apoptosis is prevented, the cells that fail to abscise can undergo subsequent attempts at cell division, resulting in severe disruption to the growing neuroepithelium. Overall, our results show how disrupted abscission significantly impacts the structure of the apical membrane, and that p53-mediated apoptosis is critical to maintain epithelial structure. This work emphasizes the importance of studying cytokinesis regulation and failures in diverse multicellular tissues in order to understand how developmental disorders and cancers arise.

It remains unclear how a midbody that fails to abscise would regress in the dense structure of a pseudostratified epithelium, and how enlarged apical endfeet form. In isolated cell lines cultured on a 2-D substrate, it was observed that after a failed abscission, sister cells would sometimes regress their midbody and merge. Other times, sister cells could remain attached by persistent cytokinetic bridges, continue to try to divide, and form a syncytium of cells connected by bridges (Gromley et al., 2003). In 3-D epithelial models, there are only a few studies examining abscission failures in an epithelium. Experiments in zebrafish demonstrated the importance of midbody positioning to establish the apical membrane and lumen in the developing Kupffer’s vesicle (KV). Disruption of abscission in the forming KV did not cause enlarged apical endfeet; rather, polarization and lumen formation failed (Rathbun et al., 2020). By contrast, in our study, we did not observe loss of polarity, and the early neural tube lumen is established.

Two other studies of abscission failures in polarized epithelia were more reminiscent of our findings, with enlarged apical surface areas. In zebrafish gastrula epithelium, when Rab25 GTPase was mutated, morphogenic activity during epiboly “tore open” persistent apical cytokinetic bridges that failed to undergo timely abscission, creating large abnormal shaped cells (Willoughby et al., 2021). Another more closely related example was recently reported using human dorsal forebrain organoids, which form spheres of polarized neurepithelium *in vitro*. They observed that mutating citron kinase (CitK), another abscission protein important for brain growth, led to enlarged apical endfeet. They speculated this was due to premature differentiation and delamination of some neuroepithelial cells, causing the remaining apical endfeet to enlarge to fill the space (Pallavicini et al., 2024). Consistent with that idea, we found premature cell cycle exit in *Cep55* KO mouse NSCs (Little et al., 2021), which would presumably induce premature delamination. However, our data herein showing that XL endfeet are multinucleate with super numerary centrosomes lead us to favor the idea that, at least in the case of the *Cep55* and *p53* dKOs, XL endfeet are primarily caused by NSCs becoming multinucleate. Taken together, these data from different model systems suggest that failures of abscission can cause different abnormal cell group morphologies dependent on whether the epithelium is two-dimensional or three-dimensional. More work is needed in different epithelial cell types and morphologies.

An important point that emerges from our work is that p53-mediated apoptosis makes the neuroepithelium robust against failures of abscission. The single knockout of Cep55(−/−) has a relatively subtle apical membrane phenotype: apical areas are enlarged, primary cilia are slightly longer, and bi-ciliation is increased, but the membrane remains polarized and intact. It can produce a brain of relatively normal form and structure, albeit small (Little et al., 2021). However, when p53-induced apoptosis is blocked, the apical membrane structure and cilia become severely disturbed. We were surprised to find some XL endfeet had a mix of mitotic and non-mitotic nuclei. This suggests that without apoptosis, more binucleate cells survive, try to divide again, and can end up with extra nuclei and centrosomes that are dyscoordinated in the cell cycle. Thus, saving cells from dying may do more harm than good for the tissue structure. These data demonstrate the importance of p53-mediated apoptosis in neuroepithelium not only as “guardian of the genome”, but guardian of epithelial structure.

How does loss of Cep55, a midbody protein, cause phenotypes in primary cilia? Our data support an indirect effect following cytokinetic abscission defects. Primary cilium assembly and disassembly is coupled to the cell division cycle: disassembly occurs in late G2 phase prior to mitosis, and assembly occurs in early G1 phase at the same time and location as midbody abscission (at the apical membrane)(Wang & Dynlacht, 2018). We found that although the percentage of ciliated cells remained unchanged, the primary cilia were slightly longer than normal in the *Cep55* KO. Cilia length correlated with apical membrane area. In addition, the number of bi- and multi-ciliated endfeet with supernumerary centrosomes was increased in the *Cep55;p53* dKO. This supports the idea that Cep55 loss affects cilia secondarily after causing abscission delays and failures. Consistent with this notion, binucleate neurons were found to have two cilia in the CikK KO mouse (Anastas et al., 2011). Another possible indirect effect of abscission dynamics on primary cilia is via midbody remnants. Midbody remnants (MBRs) are the central bulge domains that remain after abscission cuts on both flanks (Kuriyama et al., 2025). We previously found that MBRs persist for some time on the apical membranes of embryonic forebrain neuroepithelium (McNeely & Dwyer, 2020). Another group reported in cultured monolayers of dog kidney epithelial (MDCK) cells that the MBR moved towards the center of the apical membrane and promoted ciliogenesis (Casares-Arias et al., 2020 & Bernabé-Rubio et al., 2016). Since we observed an increase in MBRs on the apical membranes of the *Cep55* KO brains (Little et al., 2021), it is possible that the longer primary cilia could be due to the excess MBRs influencing ciliogenesis.

By contrast, two recent studies proposed that Cep55 has a direct role in primary cilia, though they came to opposite conclusions about its role. The first group generated a different mouse knockout of Cep55 and examined the choroid plexus epithelial cells (CPEC) and found almost a two-fold increase in cilia length compared to controls. Through further knockdown experiments in cultured dissociated human retinal pigment epithelial (RPE1) cells, they concluded that Cep55 promotes cilium disassembly (Zhang et al., 2021). The second study suggested that Cep55 promotes primary cilium growth. They observed a decrease in the percentage of ciliated NSCs in a *Cep55* KO mouse and ciliated RPE-1 cells after Cep55 knockdown (Rashidieh et al., 2021). Although we were not able to detect endogenous Cep55 localization to centrosomes or basal bodies in mouse cells using an antibody that can detect Cep55 protein in midbodies (Little et al, 2021; and data not shown), it may simply be too low or transient at basal bodies to detect with our method. Thus, more work is needed to sort out the possible direct and indirect functions of Cep55 and abscission itself in regulating primary cilia in various cell types.

## MATERIALS AND METHODS

### Mice

Mouse colonies were maintained in accordance with NIH guidelines and policies approved by the University of Virginia IACUC. The morning of the vaginal plug was considered E0.5.

Embryos were harvested by cesarean section, and littermate embryos served as controls for all experiments, with genotypes confirmed by PCR. The Cep55 allele (strain C57BL/6N*Cep55*^em1(IMPC)Tcp^), a 600 base-pair deletion encompassing all of exon 6 and flanking intronic sequence, was made as part of the KOMP2 Phase 2 project at the Toronto Center for PhenoGenomics for the Canadian Mouse Mutant Repository. It appears to be a null allele with no detectable Cep55 protein (Little et al., 2021). It was maintained on C57BL/6 and 50%/50% C57BL/6 and FVB/N background. p53 knockout (Trp53^tm1Tyj^) mice on C57BL/6 background were obtained from The Jackson Laboratory [(Jacks et al., 1994); JAX stock #002101 The Jackson Laboratory Bar Harbor, ME]. These mice were bred with 50/50% C57BL/6 and FVB/N *Cep55* knockout embryos for creation of the *Cep55;p53* mouse line. Embryos used for experiments were from 50%/50% C57BL/6 and FVB/N background, and sex of embryonic mice were not noted as sex was not a relevant biological variable for these experiments. E14.5 embryos were utilized for all experiments.

### Immunofluorescent Staining on cryosections

To collect cryosections, embryos were collected and decapitated. Heads were put in 4% PFA (paraformaldehyde) for 4-6 hours before being transferred to 30% sucrose for a few days. Heads were then embedded in OTC (Tissue-Tek, 4583) and cryosections were cut at 20µm on a Leica CM3050S cryostat and collected on Superfrost Plus slides (Fisher Scientific, 12–550-15). For IF staining, slides were warmed up to room temperature for 30-45 minutes before being blocked in 2% normal goat serum. The primary antibody was allowed to incubate overnight at 4ºC after blocking. Sections were washed the next day with PBS then secondary antibodies (1:200) were added for 30 minutes at room temperature. Following the secondary incubation, DAPI (4’,6-diamidino-2-phenylindole) was added for 10 minutes then sections were washed before being coverslipped with VectaShield fluorescent mounting medium (Vector Laboratories Inc., H-1000).

### Cortical Slab Preparation and Immunostaining

Apical slabs were prepared as previously described (Kerstin M. Janisch & Dwyer, 2016). E14.5 embryo cortices were exposed by removing the skull and fixed with 2% PFA. After primary fixation, cortices were pinched off and flipped so that the apical membrane was upright, and the slab was trimmed until the apical surface laid flat. Slabs were fixed an additional 2 minutes with 2% PFA before incubating with PBS + 0.01% Triton for 10 minutes. Slabs were blocked using 5% Normal Goat Serum and primary antibodies as well as Phalloidin 568 (Invitrogen A12380), used to label F-actin, were added and allowed to incubate at room temperature for 1-2 hours before leaving overnight at 4ºC. Slabs were washed the next day with PBS + 0.01% Triton, then twice more with PBS before the secondary antibody and DAPI was added for 1 hour at room temperature. After secondary incubation, slabs were washed with PBS before coverslipping with VectaShield fluorescent mounting medium (Vector Laboratories Inc., H-1000).

### Antibodies

Primary antibodies used in this analysis: rabbit polyclonal Arl13B (Proteintech 17711-1-AP), Mouse monoclonal Arl13B (NeuroMab 75-287),, rabbit polyclonal FOP (Proteintech 11343-1-AP).

### Imaging and Statistical Analysis

Images for figures 1–3 were taken on an inverted DeltaVision with TrueLight deconvolution microscope with softWoRx Suite 5.5 image acquisition software (Applied Precision (GE Healthcare), Issaquah, WA). Images used for figure 3G-I were taken on a Leica Thunder with Leica LAS X software. A Leica SP8 confocal microscope with LAS X software was used to collect images in figure 4. SEM images were taken on a Zeiss Sigma VP HD field emission. All image analysis was performed on ImageJ/FIJI and statistical analysis was performed using Microsoft Excel or GraphPad PRISM software. Normality and lognormality tests were used to decide which statistical test to run on data, and specific statistical tests are indicated in figure legends. Error bars in graphs 1F, H and 3D, I represent s.e.m. (standard error of the mean).

### Apical Slab Endfoot Area, Cilia, Centrosome, and Nuclei Analysis

Apical slab images for figures 1–3 were taken on an inverted DeltaVision with TrueLight deconvolution microscope with z stack step size 0.5µm. 5 images were taken at 60x magnification of each individual slab. When analyzing, an area of ~1000µm^2^ of the image was cropped then apical endfeet were counted and cilia was observed. Cilia lengths were measured as well as apical endfoot area. For figure 3G-I, images were acquired on a Leica Thunder with LAS X software with a 63X objective. Five images were taken from each slab, and images cropped to between 1000-1100µm^2^ for analysis. Nuclei analysis images used in figure 4 were taken on a Leica SP8 confocal microscope using a 63x objective and LAS X software. Nuclei and associated apical endfeet were measured and counted, and mitotic nuclei were identified by condensed chromatin labeled with DAPI. For figure 2C and 3C, XL endfeet were defined as having an area >16µm^2^, which was higher than the 99% upper confidence interval of areas of mitotic apical endfeet in control brains.

### Scanning Electron Microscopy

Heads were decapitated from E14.5 embryos and skulls were opened to expose the cortex before being fixed in 2.5% glutaraldehyde in 0.1M cacodylate buffer (Electron Microscopy Sciences

15960) with 3mM CaCl_2_ dihydrate, then stored at 4ºC for at least 1 day. After fixation, the apical membrane of the cortex was exposed by using a no.11 scalpel to cut off the top of the cortex and create a “bowl”. The cortex “bowl” was placed in a 24 well plate and incubated in 1% osmium (Electron Microscopy Sciences 19152) for 1 hour at room temperature protected from light. Samples were rinsed in dH_2_O ~10-20 times every 5 minutes before being left in dH_2_O overnight at 4ºC. An ethanol series using 30% EtOH, 50% EtOH, 70% EtOH, 95% EtOH, 100% EtOH was used to dehydrate the tissue at 4ºC. Finally, samples were dried with a few drops of HMDS drying solution (Electron Microscopy Sciences 50-243-18) overnight. Before imaging, samples were mounted to a specimen mount (Ted Pella Inc. 16111) with 9mm carbon adhesive tabs (Electron Microscopy Sciences 77825-09) and coated with 12nm of gold. Images were taken on a Zeiss Sigma VP HD field emission SEM at the UVA Advanced Microscopy Core Facility.

## ACKNOWLEDGEMENTS

This work was supported by NIH Grants R01NS076640 and R01HD102492 to N.D.D., and UVA Brain Institute Presidential Fellowship in Collaborative Neuroscience to K.S.L. We thank Jessica N. Little and the Xiaowei Lu and Ann Sutherland labs for advice and discussion. We acknowledge Hayley Dingsdale and Klaudia Filipek for related experiments and discussion, and Clarissa Q. Nassar for help with dissections. We are grateful to the Canadian Mouse Mutant Repository for use of the *Cep55* allele sperm samples. We thank the Lu lab for sharing reagents and the Hirschi lab for use of their Leica Thunder and Confocal Microscopes.

